# Spatiotemporal Control of CRISPR/Cas9 Function in Cells and Zebrafish using Light-Activated Guide RNA

**DOI:** 10.1101/831974

**Authors:** Wenyuan Zhou, Wes Brown, Anirban Bardhan, Michael Delaney, Amber S. Ilk, Randy R. Rauen, Shoeb I. Kahn, Michael Tsang, Alexander Deiters

## Abstract

We developed a new method for conditional regulation of CRISPR/Cas9 activity in mammalian cells and zebrafish embryos via photochemically activated, caged guide RNAs. Caged gRNAs are generated by substituting four nucleobases evenly distributed throughout the 5’-protospacer region with caged nucleobases during synthesis. Caging confers complete suppression of gRNA:target dsDNA hybridization and rapid restoration of CRISPR/Cas9 function upon optical activation. This tool offers simplicity and complete programmability in design, high spatiotemporal specificity in cells and zebrafish embryos, excellent off to on switching, and stability by preserving the ability to form Cas9:gRNA ribonucleoprotein complexes. caged gRNAs are novel tools for conditional control of gene editing thereby enabling the investigation of spatiotemporally complex physiological events by obtaining a better understanding of dynamic gene regulation.

## Introduction

Adapted from the prokaryotic acquired immune system, CRISPR/Cas9 has been extensively studied and meticulously developed for its advantage in efficient and precise genome editing in a customizable fashion.^1–5^ As an RNA-guided DNA endonuclease, Cas9 protein first binds to a guide RNA (gRNA), which then enables site recognition by Cas9 on the target locus through Watson-Crick base pairing between the 5’ 20 nucleotide protospacer region of the gRNA and the desired DNA sequence. Subsequent cleavage of the target locus is then carried out by the nuclease domains, HNH and RuvC, of Cas9.^6^ Recent developments of the CRISPR/Cas9 system includes broad genomic targetability enabled by Cas9 variants with PAM promiscuity,^7^ gene activation and repression,^8–9^ nucleobase editing,^10^ genomic loci imaging,^11^ and epigenetic modifications.^12^ Based on these developments, growing concerns of unwanted genomic manipulation^13^ and desire for synchronization of CRISPR/Cas9 activity with precisely orchestrated genetic networks need to be addressed.

Aiming at higher genomic editing precision by limiting the window of CRISPR/Cas9 activity as well as probing of spatiotemporally controlled gene function, researchers have endeavored to broaden the CRISPR/Cas9 toolkit for conditional control of its activity.^14–16^ Such efforts include small molecule-induced Cas9 protein activation^17–20^ or reassembly,^21^ light activation of caged Cas9,^22^ reconstitution of single-chain Cas9^23^ and split-Cas9,^24^ as well as optically controlled recruitment of transcription factors to catalytically dead Cas9 (dCas9).^25–26^ Amongst these developments, much effort was put into the regulation of the Cas9 protein to restore its function upon external stimulation. These methods inevitably require the painstaking steps of protein engineering, including the screening of mutations and split sites,^21, 24^ directed evolution,^17^ or unnatural amino acid mutagenesis.^22^ We anticipate that conditional control of chemically modified gRNA will not only circumvent the need for protein engineering, but will also provide a more easy-to-design and direct path to regulating the interaction between Cas9:gRNA ribonucleoprotein (RNP) complex and the target dsDNA. Several previous reports have shed light on this path, including using cleavable antisense-DNA as a protector for gRNA activity,^27^ ligand-dependent RNA cleavage and deprotection,^28^ ligand-dependent recruitment of transcriptional activators to dCas9,^29^ and small molecule-induced reassembly of the Cas9:gRNA complex.^30^ These designs, however, still are limited by the requirement for a third cellular component^27–28^ or reduced gRNA stability due to inability of RNP complex formation before activation.^30–32^ This is particularly important, as RNP delivery has been established as a universal approach for gene editing in different tissues and species with high efficiency and specificity, compared to alternative editing modalities.^33–36^

We henceforth introduce a photocaged gRNA design for the direct regulation of the interaction between RNP and dsDNA using light and demonstrate its application in an animal model. 6-Nitropiperonyloxymethyl (NPOM)-caged nucleobases have been successfully applied in the light-triggering of nucleic acid base-pairing in many living organisms.^37–40^ We employed NPOM-caged uridine and guanosine (Figure 1a) for application of this approach to a select set of target sequences in both mammalian cells and zebrafish embryos (Figure 1b). By replacing regular nucleobases with NPOM-caged nucleobases within the protospacer region of the gRNA, we anticipated that Cas9:gRNA:dsDNA ternary complex formation is inhibited until photolysis restores the base-pairing capability of the gRNA, while Cas9:caged gRNA interactions remain undisturbed (Figure 1c). This design is based on the rationale that the placement of NPOM-caging groups should ensure fast and complete photolysis to optically restore hybridization of an otherwise inaccessible protospacer of the gRNA. Our past experience has shown that very little background activity and excellent off → on switching upon light activation is achieved by installing one caging group every 5-6 nucleobases, evenly distributed throughout the oligonucleotide.^41–43^ We further envisioned that the Cas9:caged gRNA RNP can be pre-assembled and delivered as a complex for improved gRNA stability,^32^ facilitating application in both cultured mammalian cells as well as zebrafish embryos by lipid-mediated transfection^44^ and microinjection,^23^ respectively. Despite its synthetic challenge, we pursued a single caged gRNA because several studies have demonstrated better stability compared to the combination of crRNA (CRISPR RNA) and tracrRNA (transactivating crRNA), when complexed with Cas9 protein.^45–46^

**Figure 1.**
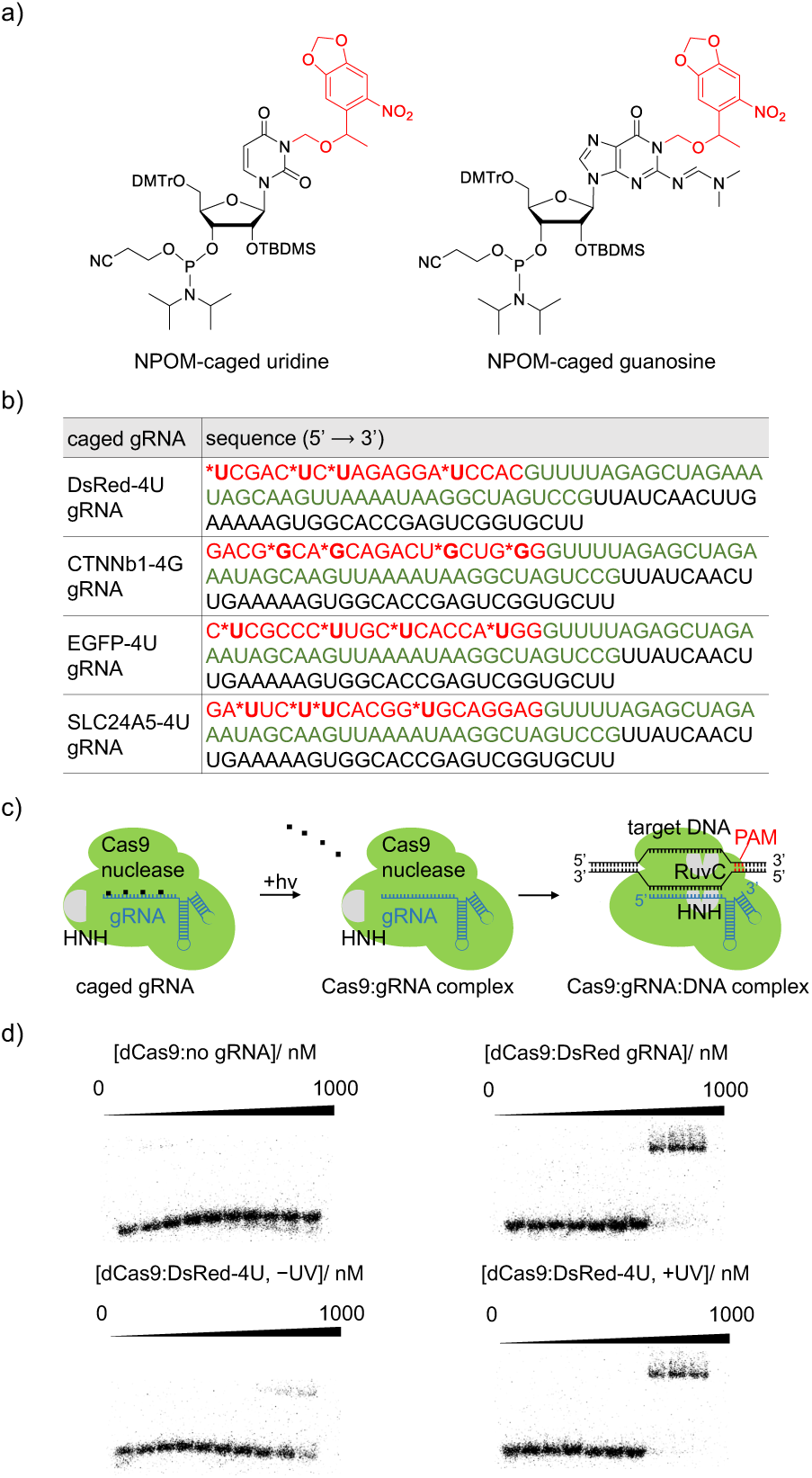
a) Structure of NPOM-protected uridine and guanosine with photocleavable caging groups shown in red. b) Sequences of photocaged gRNAs. The photocaged nucleotides are labelled by asterisks and the 20 nt base-pairing region of the gRNA is shown in red. The Cas9 binding region is shown in green and the *S. pyogenes* terminator region is shown in black. The corresponding non-caged gRNAs (DsRed, CTNNb1, EGFP, and SLC24A5) have the exact same sequences without the nucleobase caging groups. c) The NPOM-photocaging groups are designed to abolish RNP binding to the target dsDNA until they are photochemically cleaved, thereby generating an active Cas9:gRNA complex. d) Phosphorautoradiography of gel shift assays demonstrates that the photocaged gRNA abolishes the binding affinity of Cas9 to target ^32^P-labelled dsDNA and that binding is fully restored upon light activation.

## Results & Discussion

As a proof of concept, we first substituted four uridines evenly distributed within the 20 nt base-pairing region of the gRNA with photocaged uridines for complete blockade of gRNA:dsDNA hybridization.^47^ To test if caged gRNA can fully suppress base pairing and restore gRNA:dsDNA hybridization upon irradiation with UV light, gel shift assays of Cas9:gRNA complex against 32P-labelled target dsDNA were conducted with non-caged gRNA (DsRed gRNA, nomenclature is used similarly for all other genes), no gRNA, and caged gRNA (DsRed-4U gRNA) in the presence or absence of 2 min irradiation (365 nm). The binding ability of caged gRNA to dsDNA was suppressed while light-induced decaging was shown to completely restore interaction of the gRNA with the complementary dsDNA (Figure 1d). Importantly, both non-caged and caged gRNAs bind to the Cas9 protein with similar affinity, demonstrating that the caging of the protospacer region of gRNA does not interfere with formation of the Cas9:gRNA RNP complex (see Supporting Information).

Inspired by the successful optical control of the interaction between the RNP and the dsDNA, we designed photocaged gRNAs targeting different loci in both mammalian cells and zebrafish embryos following the developed strategy (Figure 1b). We first tested the optical triggering of CRISPR/Cas9 activity in mammalian cells transfected with a dual-fluorescence reporter plasmid.^48^ Targeted cleavage by Cas9 endonuclease both at the beginning and at the end of the DsRed-polyA gene cassette results in cells switching from expressing DsRed to expressing EGFP (Figure 2a). HEK293T cells harboring the dual reporter plasmid were transfected with Cas9:EGFP gRNA together with Cas9:DsRed gRNA or Cas9:DsRed-4U gRNA RNPs, and were incubated for 6 hours before irradiation with 365 nm light. It should be noted that only one caged gRNA is needed in combination with EGFP gRNA to achieve full suppression of DsRed gene excision in the absence of optical triggering and efficient editing after illumination. The cells were then incubated for 72 hours, followed by imaging. EGFP expression was only observed in the case of light exposure, indicating activation of Cas9 nuclease activity at the desired target sites by decaging of DsRed-4U gRNA while caged RNP-transfected cells that were kept in the dark remained inactive at the same minimal background level that is observed when no gRNA is present (Figure 2b). DsRed fluorescence is visible in all cells as DsRed fluorescent protein expressed before the activation of CRISPR/Cas9 is highly stable and thus is also visualized at the time of imaging. Possible insufficient editing of the transiently transfected pRG reporter could also contribute to the observed DsRed fluorescence. Quantification of the fluorescent protein expression levels was carried out by using ImageJ software. Background was first subtracted based on a fixed value determined by the fluorescence intensity of non-transfected cells.^49^ Then the fluorescence intensity of each channel for all the cells in one well was integrated to represent the expression level of the fluorescent protein (Figure 2c).^50^

**Figure 2.**
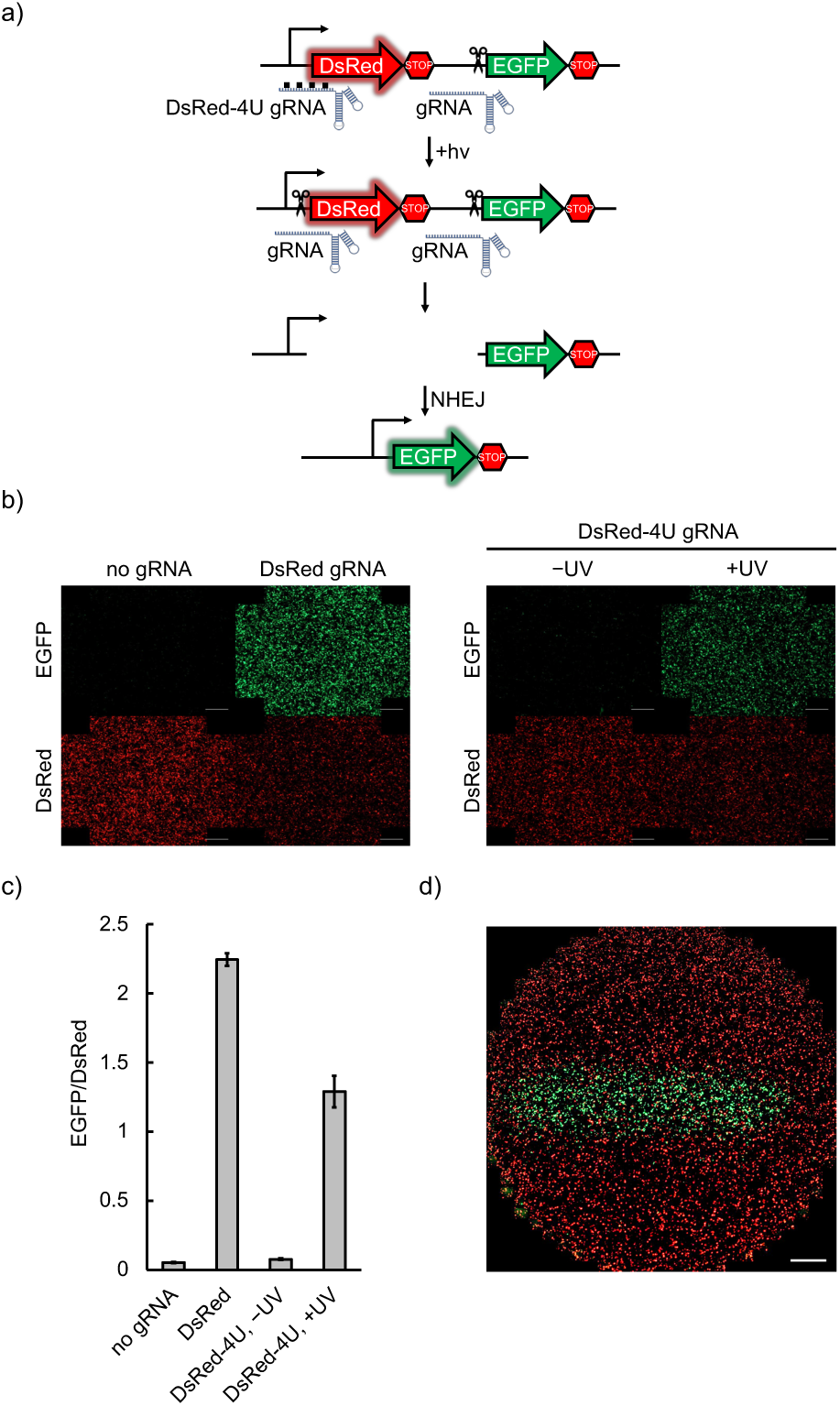
a) Schematic of the pRG reporter plasmid. Upon light activation, both functional gRNAs allow excision of the DsRed-terminator cassette from the pRG reporter plasmid and NHEJ repair leads to the expression of EGFP. b) HEK293T cells transfected with the pRG reporter plasmid, followed by delivery of Cas9:gRNA RNP complexes, were treated with or without 365 nm irradiation. EGFP expression was only observed with non-caged gRNA or when the caged gRNA was photochemically activated (scale bar = 100 μm). c) Quantification of EGFP and DsRed fluorescence was conducted by integration of fluorescence intensity in three independently transfected and treated wells for each condition using ImageJ software. d) Spatial control of light-activated Cas9:gRNA function through patterned irradiation. HEK293T cells expressing the pRG reporter plasmid and transfected with Cas9:caged gRNA complex were exposed to 365 nm irradiation through a 2 mm-wide slit in an mask (scale bar = 100 µm).

Optical control of caged gRNA presents an opportunity for precise spatial activation. Indeed, only cells exposed to 365 nm light through a slit-containing mask produced EGFP fluorescence, while all non-exposed cells only displayed DsRed expression (Figure 2d).

To demonstrate applicability of the developed optical tool to editing of the mammalian genome, we used a reported gRNA sequence (Figure 1b) to target a mammalian genomic locus within the CTNNb1 gene.^51^ Here, NPOM-caged guanosine (Figure 1a) was used instead of NPOM-caged uridine in order to achieve an even distribution of caged nucleobases following the methodology. Either CTNNb1 gRNA or caged CTNNb1-4G gRNA were delivered to HEK293T cells as Cas9 RNP complexes. Light activation was carried through exposure to 365 nm light 6 hours after delivery and gene editing was allowed for 3 days before cells were collected, followed cell lysis, and amplification of the genomic target site by nested PCR. Sanger sequencing of the amplicon and TIDE (Tracking of Indels by DEcomposition) analysis^52^ showed indel formation with 23.9% frequency for CTNNb1 gRNA and 32.2% frequency for light-activated CTNNb1-4G gRNA, while virtually no background editing was detected in the absence of irradiation of CTNNb1-4G gRNA (0.7% frequency). These results demonstrate that we are efficiently editing the mammalian cell genome with light-activated CTNNb1-4G gRNA to a similar extend as the non-caged gRNA, while no editing was observed in the absence of irradiation – showcasing the excellent off → on switching of our caged gRNA methodology.

The Cas9:gRNA RNP complex is an excellent tool for gene editing in aquatic embryos, due to ease of assembly and injection into the fertilized oocyte.^23^ Furthermore, optical control is a powerful approach for conditional gene editing in zebrafish, because the embryos are transparent during the most important stages of development, allowing for irradiation of all tissues. To demonstrate the utility of photocaged gRNAs to control Cas9 gene editing in zebrafish, we first injected RNPs assembled with a caged gRNA targeting the start codon of EGFP (EGFP-4U) in the genome of a transgenic fish line *(Tg(ubi:loxP-EGFP-loxP-mCherry)* (Figure 3a).^53^ Disruption of the start codon prevents EGFP expression, as demonstrated by representative micrographs after application of non-caged gRNA (Figure 3b). Optical activation of EGFP-4U RNP complexes in embryos had similar editing efficiency as the non-caged gRNA containing RNP, significantly reducing EGFP expression in all embryos, as determined by fluorescent imaging at 48 hours post-fertilization (hpf) (Figure 3c) and fluorescence intensity measurements (see Supporting Information). Some mosaicism can be seen at levels similar to mosaicism from RNP injection previously reported. No toxicity was observed from exposure to 365 nm light (Supporting Information).

**Figure 3.**
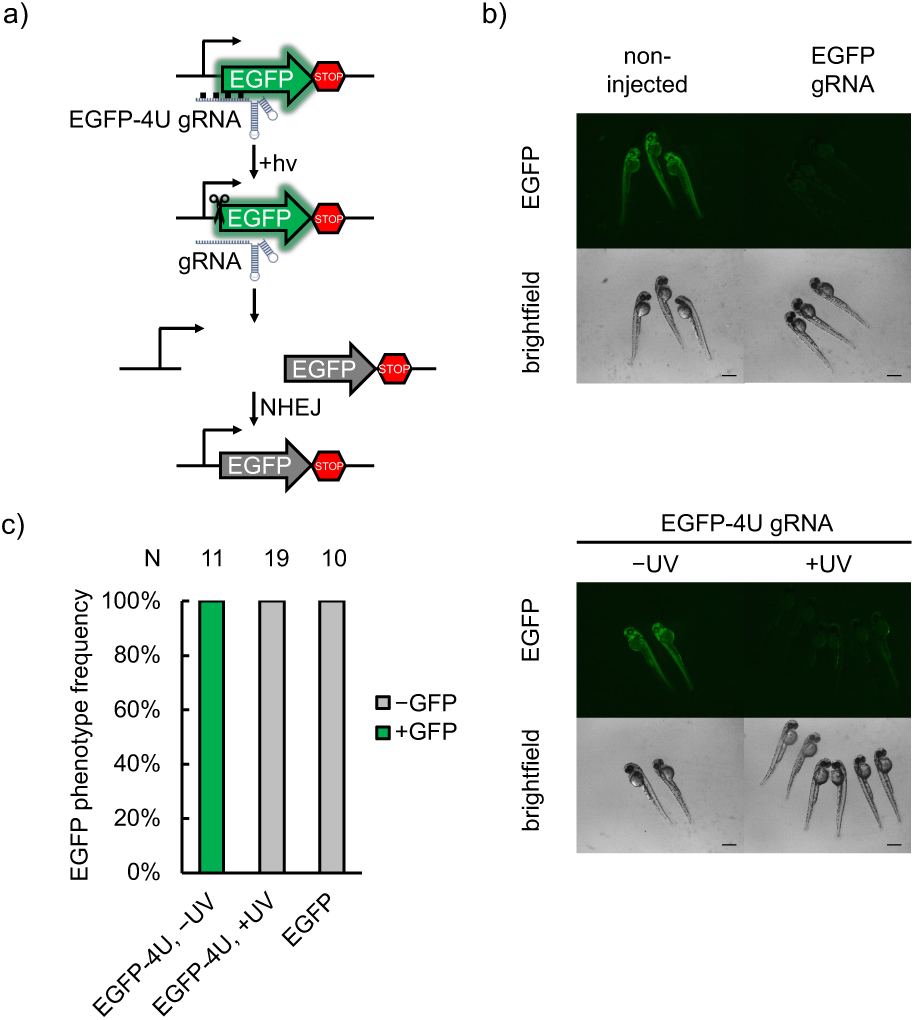
a) Schematic of the transgenic fish line fluorescent reporter. The gRNA recognizes the start codon region of EGFP, mutates it upon editing, and thus abolishes expression of the fluorescent protein. b) Representative micrographs of zebrafish at 48 hpf. Non-injected embryos demonstrate strong EGFP expression, while Cas9 RNP-injected embryos show complete loss of EGFP expression. c) Editing ability is blocked in caged EGFP-4U gRNA until irradiation with 365 nm light at 1 hour post-injection, indicating optical control of gene editing in embryos. Phenotype frequencies of the injected embryos at 48 hpf are shown. Scale bars = 300 µm.

In order to further validate the universal applicability of this methodology, we next targeted the *slc24a5* gene,^54^ an endogenous gene in zebrafish which is important for development of pigmentation by 48 hpf. *slc24a5* has been edited with Cas9 RNP injection before,^55–56^ and the lack of pigmentation induced by editing of this locus is commonly referred to as the golden phenotype, most robustly observed as pigment loss in the retina of the developing animal. We used the same gRNA sequence that has been previously used for disrupting *slc24a5* function.^56^ Four uridine bases in the protospacer region were replaced with NPOM-caged uridines and showed complete inhibition of Cas9 editing function until it was restored upon irradiation with 365 nm light. Resulting gene editing led to almost complete loss of retinal pigmentation (Figure 4a). Increasing light exposure (5 min vs 3 min) resulted in increased pigmentation loss in a larger proportion of embryos, demonstrating that this method allows for tuning of editing efficiency and that full optical activation of Cas9 RNP function can be achieved (Figure 4b, and Supporting Information). Optical off to on switching of gene editing is further confirmed by TIDE analysis of the *slc24a5* gene locus, showing a 75% indel rate for non-caged SLC24A5 gRNA, 84% for light-triggered SLC24A5-4U gRNA, and 7% for SLC24A5-4U gRNA in the absence of irradiation.

**Figure 4.**
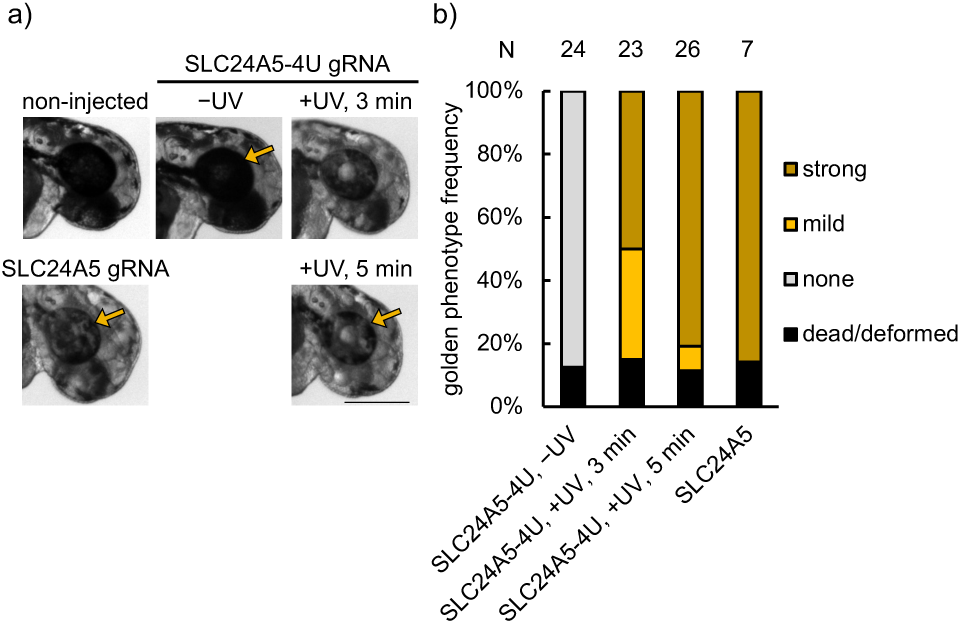
a) Representative images of embryos injected with RNPs assembled from non-caged SLC24A5 gRNA or caged SLC24A5-4U gRNA, then non-irradiated or irradiated (365 nm) for 3 or 5 min. Images were recorded at 48 hpf. Arrows point to the retina, demonstrating loss of pigmentation through successful editing of the SLC24A5 locus. Scale bar = 300 µm. b) Phenotype frequencies of the injected embryos at 48 hpf. The golden phenotype was measured based on retinal pigmentation, with mild representing small patches of pigment loss, and strong representing a majority or complete retinal pigment loss. No background activity was observed for SLC24A5-4U, while 365 nm exposure for 5 min activated gene editing to the same level as the non-caged gRNA.

## Summary

We developed a new method to optically control CRISPR/Cas9 activity through nucleobase-caged gRNAs, thereby further expanding the light-activated tool set available for conditionally controlled gene editing with spatial and temporal resolution.^57–59^ We successfully applied this approach in both mammalian cells and zebrafish embryos with high efficiency on both transiently transfected plasmid DNA and genomic loci. NPOM-caged gRNAs expand the gene editing toolbox with unique features, including 1) rapid and non-invasive activation of CRISPR/Cas9 activity, 2) precise spatiotemporal control, 3) modularity and programmability of the light-controlled gRNA sequence, 4) formation of a stable Cas9:gRNA complex from commercially available protein, 5) broad applicability for delivery into cells and organisms in the form of RNP complexes, and 6) capability for tuning of gene editing efficiency. We expect that NPOM-caged gRNAs will find utility in the dissection of regulatory networks in the rapidly developing zebrafish embryo in a temporally and spatially sensitive manner. Furthermore, the light-activated Cas9:gRNA RNP system is expected to be functional in other cell lines and (aquatic) embryos.

## Acknowledgements

This work was supported by the National Institutes of Health (R21HD085206 and R01GM112728 to AD; T32GM088119 to WB).

