## Supporting Information for "Spatiotemporal Control of CRISPR/Cas9 Function in Cells and Zebrafish using Light-Activated Guide RNA"

### Methods

#### Caged gRNA syntheses

The NPOM-caged uridine and guanosine phosphoroamidites were synthesized according to established protocols by Dharmacon, Horizon Discovery, Inc.[1]

Caged and non-caged oligonucleotides were synthesized by Dharmacon using an LGC Bioautomation MerMade 4 DNA/RNA Synthesizer (Irving, TX, USA) employing standard β-cyanoethyl phosphoramidite chemistry. The caged gRNAs were synthesized at 1-μmole scale on 1000 Å rU derivatized CPG solid phase supports obtained from LGC Biosearch Technologies (Novato, CA, USA) using 5'-DMTr-ON approach. Reagents and non-caged phosphoramidites for automated RNA synthesis were also obtained from LGC Bioautomation. Optimized synthesis cycles with coupling time of 8 mins were employed for both non-caged and caged phosphoramidites at 0.08 M concentration. Coupling efficiency in each cycle was monitored by following the release of dimethoxytrityl (DMTr) cations after each deprotection step. No significant loss of DMTr was noted following the addition of caged-U or caged-G phosphoramidites to the oligonucleotide.

Following synthesis, cleavage from support and deprotection of the oligonucleotides were carried out with 1 mL of AMA (ammonium hydroxide/40% aq. methylamine 1:1 v/v) solution at 65 °C for 20 min. The TBDMS group (*tert*-butyldimethylsilyl) was deprotected using 65 μL of TEA (triethylamine) and 75 μL of TEA-3HF (triethylamine trihydrofluoride) in 110 μL of DMSO at 65 °C for 2 h.

#### Caged gRNA purification

Purification was performed on a Waters Autopurification system using a Waters X-Bridge BEH C18 column (10 mm x 100 mm, 130 Å, 5 μm). The buffers used were 50 mM TEAA (triethylammonium acetate) with 5% MeOH (methanol) as buffer A and 100% MeOH as buffer B. The oligonucleotides were purified using a gradient of 15-60% buffer B over 60 mins at a flow rate of 4 mL/min. Fractions were analyzed by UPLC and the highest purity fractions were pooled together.

Final deprotection of the 5'-DMTr group was accomplished by treating the purified pool with 400 μL of 35 mM NaOAc (sodium acetate) at 55 °C for 2 h. The oligonucleotides were then precipitated using 50 μL of 3 M NaOAc and 1.5 mL of cold ethanol. Following incubation at -80 °C for 30 mins, the oligonucleotides were pelleted by centrifugation, quantified at 260 nm and further analyzed by UPLC and ESI-LCMS.

#### UPLC and ESI-LCMS analyses

The oligonucleotides were analyzed on a Waters Aquity UPLC system using a Waters BEH C18 column (2.1 mm x 50 mm x 1.7 μm). Samples were prepared by dissolving 0.5 nmol of the oligonucleotide in 75 μL of water and injecting 2 μL of the solution. The buffers used were 50 mM Dimethylhexylammonium acetate with 10% $CH_3CN$ (acetonitrile) as buffer A and 50 mM Dimethylhexylammonium acetate with 75% $CH_3CN$. The gradient utilized for analysis was 25-75% buffer B over 5 mins with a flow rate of 0.5 mL/min at 60 °C.

ESI-LCMS data for the oligonucleotides were acquired on a Thermo Ultimate 3000-LTQ-XL mass spectrophotometer. Samples were prepared by dissolving 0.5 nmol of the oligonucleotide in 75 μL of water and injecting 10 μL of the solution using a Novatia C18 (HTCS-HTC1-4) trap column. Following injection, sample was eluted into the LTQ-MS with 85% $CH_3CN$, 50 mM HFIP (hexafluoro-2-propanol), 10 μM EDTA (ethylenediaminetetraacetic acid), 0.35% DIPEA (N,N-diisopropylethylamine) and analyzed for mass (**Supporting Figures 1-4**).

#### Non-caged gRNA preparation

Template sequences for *in vitro* transcription of gRNAs were listed in **Table 1**. Transcription templates were prepared by mixing 10 µg of the Forward strand, 10 µg of the Reverse strand, 1 µL of 10x TE/$Mg^{2+}$ buffer, and nuclease free water to make up a 10 µL annealing mixture, followed by heating at 95 °C for 3 min and cooling down to 25 °C at a rate of −0.2 °C/sec, using a thermal cycler (Bio-Rad, T100). The T7 or SP6 MEGAscript kits (Fisher) were used for *in vitro* transcription of non-caged gRNAs. In a 20 µL transcription mixture, 1 µg of the annealed double strand template was used following manufacturer's protocol, for 16 hrs at 37 °C. The transcription was stopped by adding 1 µL of Turbo DNase from the MEGAscript kit and kept at 37 °C for 15 min. For extraction of the transcribed gRNA, the reaction mixture was mixed with 15 µL Ammonium Acetate Stop Solution (5 M ammonium acetate, 100 mM EDTA), 115 µL TRIzol reagent (Fisher), and 300 µL chloroform (Fisher). After brief vortexing, the mixture was centrifuged at 13,200 rpm at 4 °C for 10 min, after which the top aqueous layer was transferred into 500 µL 2-propanol (Fisher). After brief vortexing, the tube was frozen at −80 °C for at least 2 hours. The tube was then centrifuged at 13,200 rpm at 4 °C for 20 min, after which the pellet was resuspended in 20 µL of nuclease-free water. The concentration of each non-caged gRNA solution was determined by absorption at 260 nm using a NanoDrop spectrophotometer and the purity was examined by 10% 6 M urea denaturing TBE PAGE (25 W, 30 min), followed by SYBR Gold (Invitrogen) staining at room temperature for 30 min (1:10000 dilution in 1x TBE buffer) and imaging using the Bio-Rad ChemiDoc™ MP Imaging System (**Supporting Figure 5**).

**Table 1.** Oligonucleotide sequences.

| Name | Sequence (5'-3') |
|---|---|
| T7-DsRed_F (pRG) | TAATACGACTCACTATAGGGAGATCGACTCTAGAGGATCCACG<br>TTTTAGAGCTAGAAATAGCAAGTTAAAATAAGGCTAGTCCGTTA<br>TCAACTTGAAAAAGTGGCACCGAGTCGGTGCTT |
| T7-DsRed_R (pRG) | AAGCACCGACTCGGTGCCACTTTTTCAAGTTGATAACGGACTA<br>GCCTTATTTTAACTTGCTATTTCTAGCTCTAAAACGTGGATCCTC<br>TAGAGTCGATCTCCCTATAGTGAGTCGTATTA |
| T7-EGFP_F (pRG) | TAATACGACTCACTATAGGGAGATAGCTAGTCTAGGTCGATGC<br>GTTTTAGAGCTAGAAATAGCAAGTTAAAATAAGGCTAGTCCGTT<br>ATCAACTTGAAAAAGTGGCACCGAGTCGGTGCTT |
| T7-EGFP_R (pRG) | AAGCACCGACTCGGTGCCACTTTTTCAAGTTGATAACGGACTA<br>GCCTTATTTTAACTTGCTATTTCTAGCTCTAAAACGCATCGACCT<br>AGACTAGCTATCTCCCTATAGTGAGTCGTATTA |
| SP6-CTNNb1_F (mammalian) | ATTTAGGTGACACTATAGACGGCAGCAGACTGCTGGGGTTTTA<br>GAGCTAGAAATAGCAAGTTAAAATAAGGCTAGTCCGTTATCAAC<br>TTGAAAAAGTGGCACCGAGTCGGTGCTTTT |
| SP6-CTNNb1_R (mammalian) | AAAAGCACCGACTCGGTGCCACTTTTTCAAGTTGATAACGGACT<br>AGCCTTATTTTAACTTGCTATTTCTAGCTCTAAAACCCCAGCAGT<br>CTGCTGCCGTCTATAGTGTCACCTAAAT |
| SP6-EGFP_F (zebrafish) | ATTTAGGTGACACTATAGCTCGCCCTTGCTCACCATGGGTTTTA<br>GAGCTAGAAATAGCAAGTTAAAATAAGGCTAGTCCGTTATCAAC<br>TTGAAAAAGTGGCACCGAGTCGGTGCTT |
| SP6-EGFP_R (zebrafish) | AAGCACCGACTCGGTGCCACTTTTTCAAGTTGATAACGGACTA<br>GCCTTATTTTAACTTGCTATTTCTAGCTCTAAAACCCATGGTGA<br>GCAAGGGCGAGCTATAGTGTCACCTAAAT |

**Table 1**. cont.

| | |
|---|---|
| SP6-SLC24A5_F (zebrafish) | ATTTAGGTGACACTATAGATTCTTCACGGTGCAGGAGGTTTTAGAGCTAGAAATAGCAAGTTAAAATAAGGCTAGTCCGTTATCAACTTGAAAAAGTGGCACCGAGTCGGTGCTTTT |
| SP6-SLC24A5_R (zebrafish) | AAAAGCACCGACTCGGTGCCACTTTTTCAAGTTGATAACGGACTAGCCTTATTTTAACTTGCTATTTCTAGCTCTAAAACCTCCTGCACCGTGAAGAATCTATAGTGTCACCTAAAT |
| Cas9 D10A_R | GTGCCGATCGCTAAGCCTATTGAGTATTTCTTATCC |
| Cas9 H840A_F | GATGTCGATGCGATTGTTCCACAAAGTTTCCTTAAAGACG |
| Cas9 H840A_R | GTGGAACAATCGCATCGACATCATAATCACTTAAACG |
| pBAD_Cas9_F | CCCATGGATAAGAAATACTCAATAG |
| pBAD_Cas9-6xHis_R | GCGCGCCGTTTAAACAAAGCTTTAATGATGATGATGATGATGGTCACCTCCTAGCTGAC |
| DsRed 55mer target_F [b)] | CGCAAGCTTCCTAGACTAGTCGACTCTAGAGGATCCACCGGTCGCCACCATGGCC |
| DsRed 55mer target_R [b)] | GGCCATGGTGGCGACCGGTGGATCCTCTAGAGTCGACTAGTCTAGGAAGCTTGCG |
| CTNNb1_F1 [c)] | GATCATACTTGTTGCAGCTTCGACAA |
| CTNNb1_F2 [c)] | CTAGTGACAAGTGGAACCAGAT |
| CTNNb1_R2 [c)] | CTACGAAGTTTGGCTCCGAGA |
| CTNNb1_R1 [c)] | CCCGCTGCACTTAGAGTTTAGTTG |
| SLC24A5_F1 [c)] | CTCTGTTACTGTCAACTCATTGTGTATTAT |
| SLC24A5_R1 [c)] | CACTGACGGATCTCTGCACT |
| SLC24A5_F2 [c)] | GCAGTTCTGAAATGATTTGTGTGTGT |
| SLC24A5_R2 [c)] | ACGAGCTCTGGAGCCGAACTC |

[a)] T7 and SP6 promoters are underlined. [b)] 55-mer dsDNA strands used for gel shift assay. [c)] Primers used in nested PCR.

##### Expression and purification of catalytically dead Cas9

The *S. pyogenes* Cas9 (abbreviated Cas9) gene was a kind gift from the Asokan Lab (North Carolina State University). D10A and H840A mutations were introduced by site-directed mutagenesis with primers mentioned in **Table 1** to render it catalytically inactive (dead Cas9, or dCas9) (**Table 1**). dCas9 was then amplified by the pBAD_Cas9_F and pBAD_Cas9-6xHis_R primers (**Table 1**), digested by NcoI and PmeI restriction enzymes (NEB, 37 °C, overnight), and purified by PCR Cleanup kit (Omega Biotek). The pBAD plasmid was digested by NcoI and PmeI restriction enzymes (NEB, 37 °C, overnight), and purified by 0.8% agarose gel and DNA Gel Extraction Kit (Omega Biotek). T4 ligation of the insert and the vector (1:3 molar ratio) was carried out at 16 °C, overnight, with 1 µL 10x T4 Ligase Buffer and 0.5 µL T4 Ligase (NEB) in a 10 µL reaction mixture. Successful site-directed mutagenesis and construction of the pBAD_dCas9-6xHis expression plasmid were confirmed by Sanger sequencing (Genewiz, USA). The plasmid map is shown in **Supporting Figure 6a.**

pBAD_dCas9-6xHis was transformed by heat shock in TOP10 *E. coli* competent cells. Overnight starter culture (containing 12 μg/mL tetracycline) was diluted 1:100 into 25 mL of LB broth (containing 12 μg/mL tetracycline) and let grow until O.D. 600 reaches 0.6, at which point, the cells were induced by 0.2% L-arabinose and protein expression went on at 16 °C for 16 hours. After 16 hours, the cells were harvested, resuspended in 5 mL lysis buffer (20 mM Tris-HCl, 200 mM NaCl, and 10 mM imidazole, pH 8.0), lysed by 1 mg/mL lysozyme (Fisher) and sonification (Fisher Sonic Dismembrator 550, 15% amplitude, 10 sec ON, 10 sec OFF, 5 min total) in the presence of protease inhibitor cocktail (Sigma) at 4 °C. His-tagged dCas9 was then pulled-down by 80 μL Ni-NTA agarose resin (G-Biosciences), washed twice by 500 μL washing buffer (20 mM Tris-HCl, 200 mM NaCl, and 50 mM imidazole, pH 8.0), eluted by 300 μL elution buffer (20 mM Tris-HCl, 200 mM NaCl, and 300 mM imidazole, pH 8.0), and dialyzed against 1 L Cas9 dialysis buffer (20 mM HEPES pH 7.5, 150 mM KCl, 1 mM TCEP, 10% glycerol) at 4 °C overnight.[2] Dialyzed dCas9 protein was then concentrated to ~ 3 uM using 50 kDa spin columns (Millipore) at 4 °C at 14000 *g* for 20 min.

The purity and concentration of dCas9-6xHis was analyzed by 6% SDS-PAGE (**Supporting Figure 7**). The gel was photographed by the Bio-Rad ChemiDoc™ MP Imaging System and the quantification was done by integrating the pixels of each band using the ImageLab 5.0 software.

**Gel shift assay for dsDNA binding and gRNA binding**

Gel shift assay was carried out following a published method, with a $^{32}$P-labelled dsDNA target.[3] For the isotopic labelling of the target dsDNA, the following protocol was used. The forward strand ssDNA (25 pmol) was mixed with 2 μL 10x T4 PNK buffer (NEB), 1.5 μL gamma-$^{32}$P-ATP (PerkinElmer, 3000 Ci/mmol), 0.8 μL T4 PNK (NEB), and nuclease-free water to make a 20 μL labeling mixture, which was incubated at 37 °C for 4 hrs, followed by heat inactivation at 65 °C for 20 min. The isotopically labelled forward ssDNA (25 pmol) was then mixed with 10x excess of reverse ssDNA (250 pmol), 10xTE/$Mg^{2+}$ buffer, and nuclease-free water to make a 50 μL annealing mixture. Annealing was carried out by heating at 95 °C for 3 min and cooling down to 25 °C at a rate of −0.2 °C/sec. The mixture was then used as a 100 nM $^{32}$P-dsDNA target solution in the gel shift assay.

Purified dCas9-6xHis (abbreviated as dCas9) was mixed with 10x molar excess of the designated gRNA and incubated at 37 °C for 10 min. Then varying concentrations of the Cas9:gRNA complex (0-1000 nM) was mixed with 1 nM of $^{32}$P-dsDNA and incubated at 37 °C for 1 hr. Afterwards, the mixtures were run on 5% native PAGE gel (in 0.5x TBE + 5 mM $MgCl_2$ running buffer) at 100 V and 4 °C for 2 hrs. Then the native PAGE gels were exposed to a phosphor screen for at least 4 hours and visualized by Typhoon FLA7000 IP Phosphoimager (GE Healthcare).

The labeling of non-caged DsRed gRNA and DsRed-4U gRNA were carried out following a published method.[4] In short, the gRNAs were diluted to 0.67 nM, mixed with 0-1000 nM dCas9, and incubated at 37 °C for 1 hr. The mixtures were then run on 5% native TBE PAGE as mentioned above, stained by SYBR Gold (1:10000) for 30 min, and visualized by the Bio-Rad ChemiDoc™ MP Imaging System (**Supporting Figure 8**).

**Mammalian cell culture, transfection, and Cas9:gRNA RNP delivery**

HEK293T cells were acquired from ATCC and maintained in DMEM media (Fisher) with 10% FBS (Fisher/USB) without antibiotics. Cells were sequentially transfected twice for the pRG reporter plasmid (**Supporting Figure 6b**) and the Cas9:gRNA RNP. For the first transfection of the pIRG reporter plasmid, 500 ng DNA was combined with 3.23 μg of linear polyethylenimine (Polysciences) per well of a 12-well plate. Six hours after the transfection of the reporter plasmid, cells were gently lifted with 100 μL TrypLE Express (Gibco), spun down at 1000 rpm at room temperature for 5 min, resuspended in fresh DMEM with 10% FBS (no antibiotics), reverse transfected with Cas9:RNP complexes, and left to grow in 96-well plates at 37 °C, 5% CO2. The

RNP complex was pre-assembled by mixing EnGenCas9 (dual-NLS-Cas9, New England Biolabs) and gRNA at a 1:1 molar ratio and incubated at 37 °C for 10 minutes. For reverse transfection of one well in a 96-well format, 1.5 pmol total RNP was mixed with 0.75 uL Lipofectamine 3000 (Fisher) in 20 µL Opti-MEM (Gibco) for 10 minutes at room temperature before mixing with 80 µL of HEK293T cells ($5x10^4$ cells) transfected with pRG dual-fluorescence reporter. For editing of the CTNNb1 gene, transfection of the pRG plasmid was excluded. HEK293T cells were only reverse transfected once with the corresponding RNP complexes.

**Light activation of RNP complexes delivered into mammalian cells**

HEK293T cells transfected with Cas9:gRNA RNP complexes were allowed 6 hours to attach to the bottom of the plate. Afterwards, fresh DMEM replaces the transfection mixture and the cells were subject to 5 minutes of 365 nm irradiation by a UV transilluminator (25 W). Fluorescent imaging of the cells was performed after 72 hours post irradiation. Media was replaced with clear DMEM-high modified growth media (Thermo Scientific) for microscopy imaging on a Zeiss Observer Z1 microscope (10X objective, NA 0.8 plan-apochromat) with EGFP (38 HE; ex: BP470/40; em: BP525/50) and DsRed (43 HE; ex: BP550/25; em: BP605/70) filter cubes, then processed and quantified in Zen 2 imaging software. For the spatial control experiments, UV irradiations were performed through a tin foil mask to only expose a subset of cells to 365 nm light for 5 minutes. Microscopy imaging was then performed in a tiled grid and stitched using the Zen 2 software. Quantification was carried by first setting a lower threshold with a fixed value to subtract background, which was determined by the fluorescence intensity of non-transfected HEK293T cells in the same plate. Then the fluorescence intensity for each channel was integrated to represent the expression level of the corresponding fluorescent protein. Editing efficiency was calculated by dividing integrated EGFP fluorescence by integrated DsRed fluorescence of each well and error bars represent standard deviation of three independently transfected and treated wells in parallel.

**Genomic PCR and TIDE analysis**

HEK293T cells were lysed by heating cell suspension at 95 °C (in 1x PBS) for 10min. Nested PCR was then carried out with cell lysates using the corresponding primers (**Table 1**). Zebrafish embryos were lysed following a published protocol.[5] In short, whole embryos were transferred to 20 µL lysis buffer (50 mM KCl, 10 mM Tris-HCl pH 8.3, 0.3% Tween 20, 0.3% NP40. Add fresh Proteinase K to a final concentration of 1 mg/mL on the day of use) and incubated at 55 °C for 16 hrs. Then the mixture was heated to 95 °C for 15 min to inactivate Proteinase K. Nested PCR was then carried out with zebrafish embryo lysates with the corresponding primers (**Table 1**).

Amplified PCR products were purified by PCR Cleanup kit (Omega Biotek) and sent to Genewiz, USA for sequencing. TIDE analysis was carried out by an online tool (https://tide.deskgen.com/). Sanger sequencing chromatograms and TIDE analyses results are shown in **Supporting Figure 9** and **Supporting Figure 10**.

**Zebrafish embryo injection and irradiation**

The zebrafish experiments were performed according to a protocol approved by the Institutional Animal Care and Use Committee (IACUC) at the University of Pittsburgh. Embryos were collected after natural mating of AB* or *Tg(ubi:loxP-EGFP-loxP-mCherry)*[6] fish lines. This transgenic fish line was a kind gift from the Leonard Zon lab (Harvard Medical School). An injection solution of 10 µM EnGen® Spy Cas9 NLS (dual-NLS Cas9, NEB) and 10 µM gRNA were incubated for 20 minutes at 37 °C. Phenol red was added to a final concentration of 0.05% as a tracer for injection. EGFP-RNP (1 nL) or SLC24A5-RNP (2 nL) was injected underneath the cell within the yolk of the single-cell stage embryo using a World Precision Instruments Pneumatic PicoPump injector. Within the first hour after injection, embryos were irradiated with 365 nm light using a handheld transilluminator placed on top of a 35 mm petri dish containing the embryos suspended in E3 water. The embryos were then incubated at 28.5 °C until 48 hpf when they were assessed for phenotype. Toxicity from UV irradiation was analyzed at 48 hpf after irradiation of non-injected embryos for 3 or 5 minutes (**Supporting Figure 11**) and no detrimental effects were observed. At 48 hpf, the embryos were imaged with a Leica M205 FA microscope with EGFP filter cube (ex: 470/40 nm, em: 525/50 nm) and the bright field channel (**Supporting Figure 12** and **Supporting Figure 13**). Fluorescence intensity was measured using ImageJ software.

#### Supporting Figures

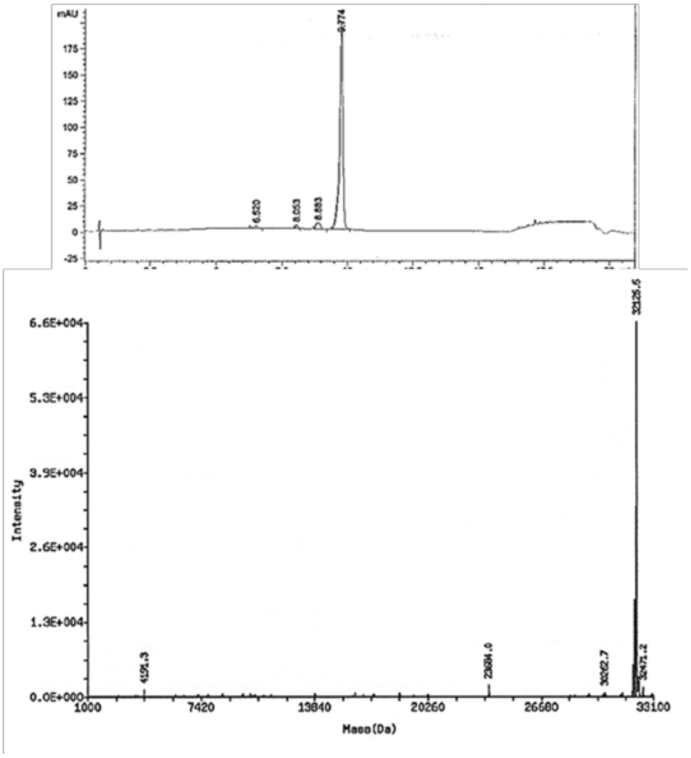

**Supporting Figure 1.** LC chromatogram and ESI-LCMS spectrum of DsRed-4U gRNA (pRG). Calculated mass: 32124 Da; observed mass: 32127 Da.

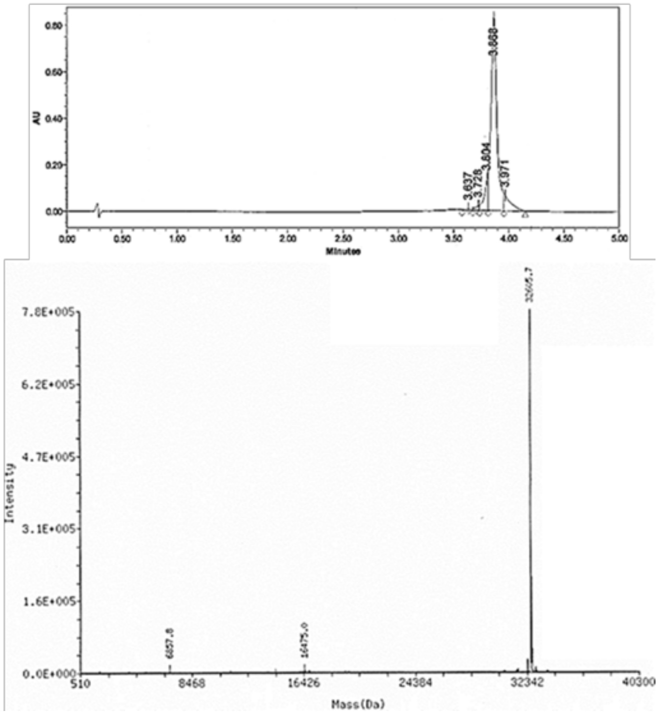

**Supporting Figure 2.** LC chromatogram and ESI-LCMS spectrum of CTNNb1-4G gRNA. Calculated mass: 32601 Da; observed mass: 32606 Da.

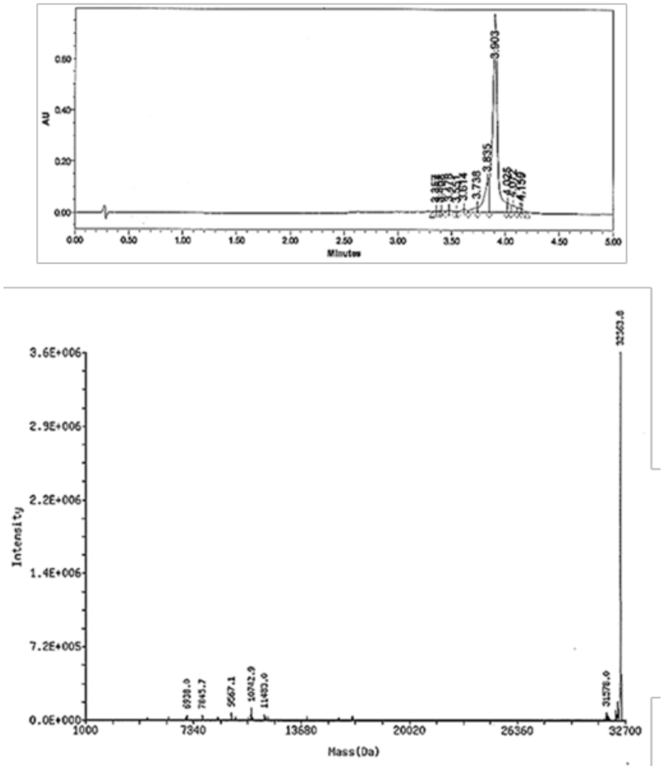

**Supporting Figure 3.** LC chromatogram and ESI-LCMS spectrum of EGFP-4U gRNA (zebrafish). Calculated mass: 32356 Da; observed mass: 32364 Da.

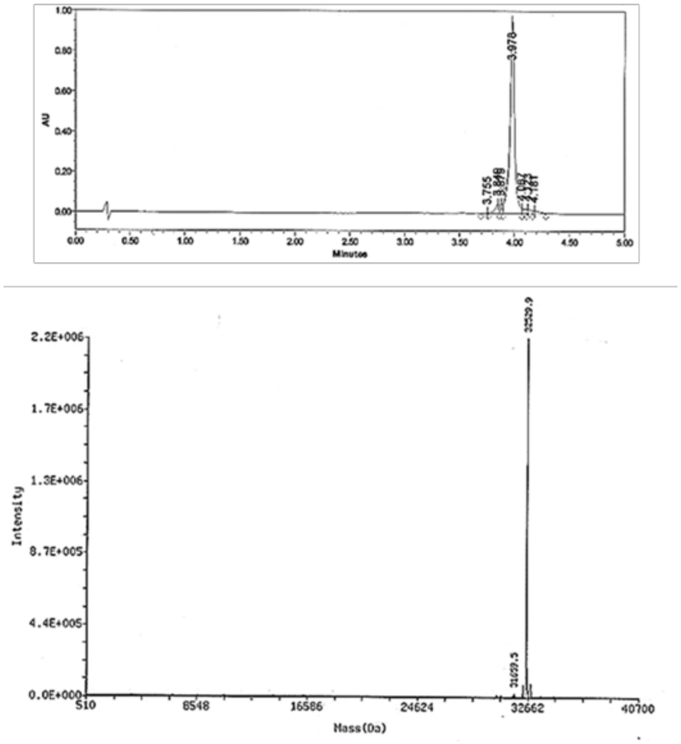

**Supporting Figure 4.** LC chromatogram and ESI-LCMS spectrum of SLC45A-4U gRNA. Calculated mass: 32524 Da; observed mass: 32530 Da.

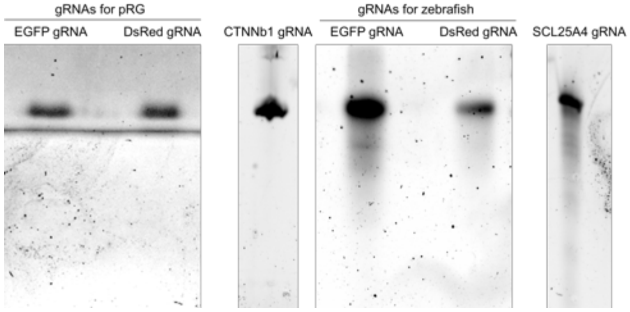

**Supporting Figure 5.** Denaturing 6 M urea TBE PAGE for all the *in vitro* transcribed, non-caged gRNAs.

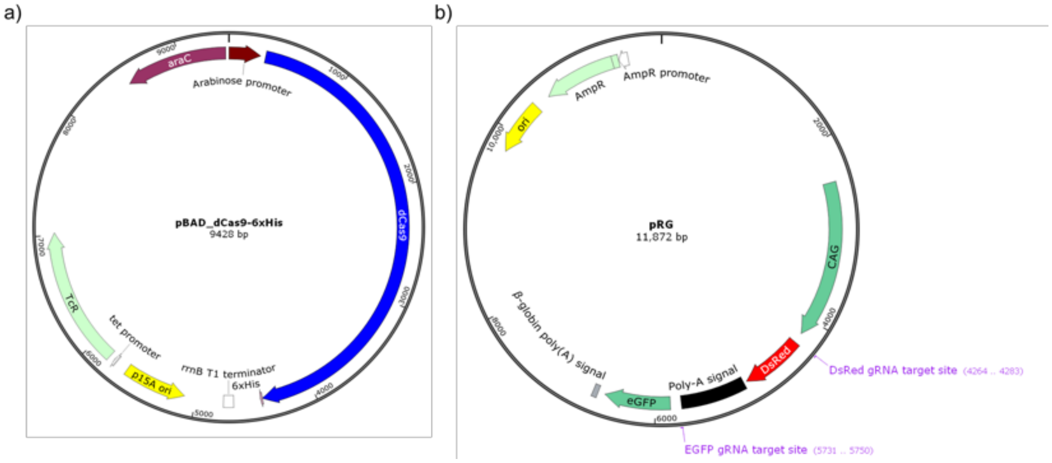

**Supporting Figure 6.** Plasmid maps of the plasmids used in this research (generate by SnapGene). a) Plasmid map for pBAD_dCas9-6xHis. b) Plasmid map for the pRG dual-fluorescence reporter plasmid.

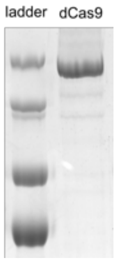

**Supporting Figure 7.** Coomassie stained SDS-PAGE analysis of dCas9-6xHis with Precision Plus Protein™ Dual Color Standards (Bio-Rad).

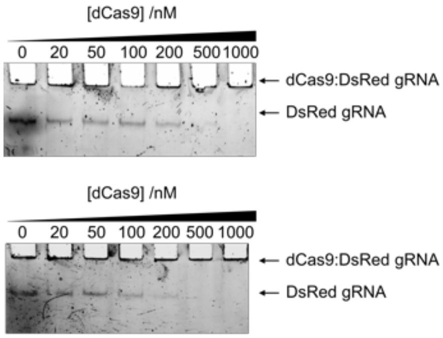

**Supporting Figure 8.** Gel shift assay of dCas9 with non-caged DsRed gRNA or DsRed-4U gRNA shows that photocaging the protospacer region of gRNA does not alter its binding to Cas9.

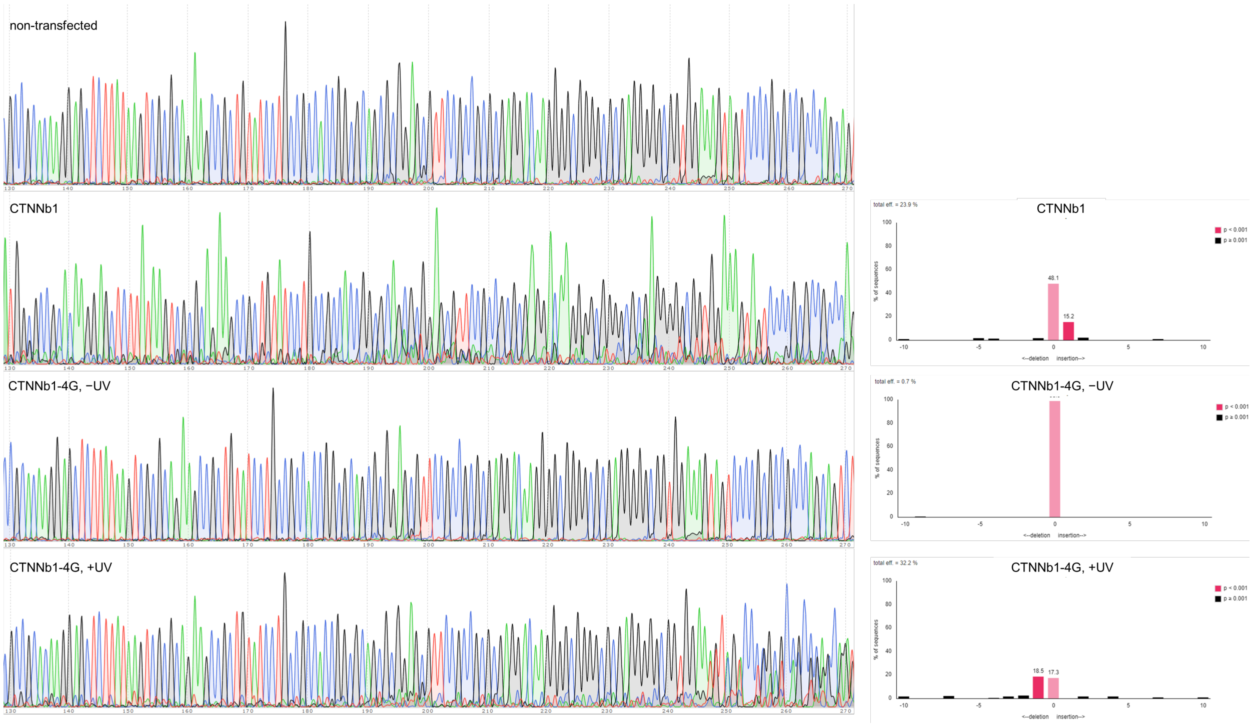

**Supporting Figure 9.** Sanger sequencing chromatograms (left) and TIDE analyses results (right) for genomic PCR products from non-transfected HEK293T cells as well as HEK293T cells transfected with non-caged CTNNb1 gRNA or caged CTNNb1 gRNA (-/+ UV) RNPs. Indels are reflected by the additional signals compared to non-transfected cells.

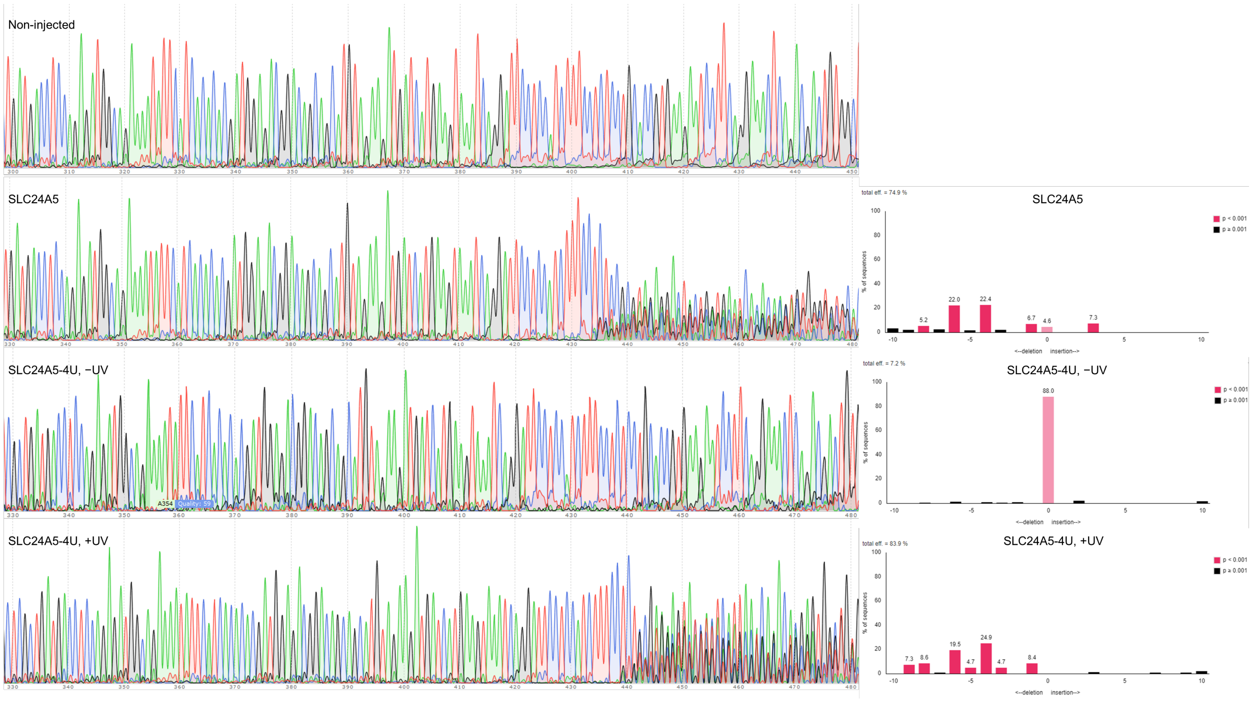

**Supporting Figure 10.** Sanger sequencing chromatograms (left) and TIDE analyses results (right) for genomic PCR products from the non-injected zebrafish embryo as well as embryos injected with non-caged SLC24A5 gRNA or caged SLC24A5 gRNA (-/+ UV) RNPs. Indels are reflected by the additional signals compared to the non-injected embryo.

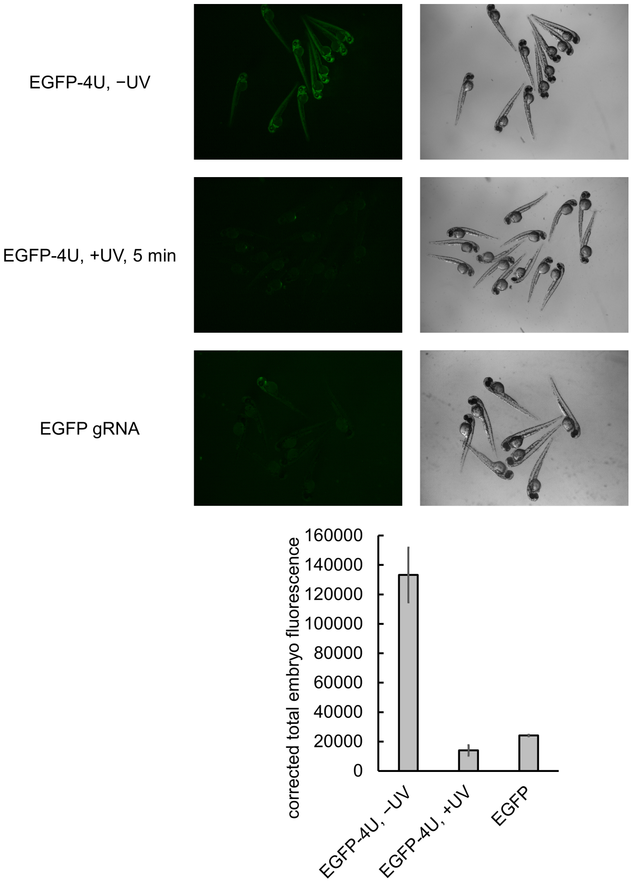

**Supporting Figure 11.** Micrographs of embryos injected with RNP targeting EGFP and irradiated one hour after injection, then imaged at 48 hpf. Fluorescence intensity in the EGFP channel of three embryos in each condition was measured using ImageJ software and presented in the chart below. Bars represent means and error bars represent standard deviations.

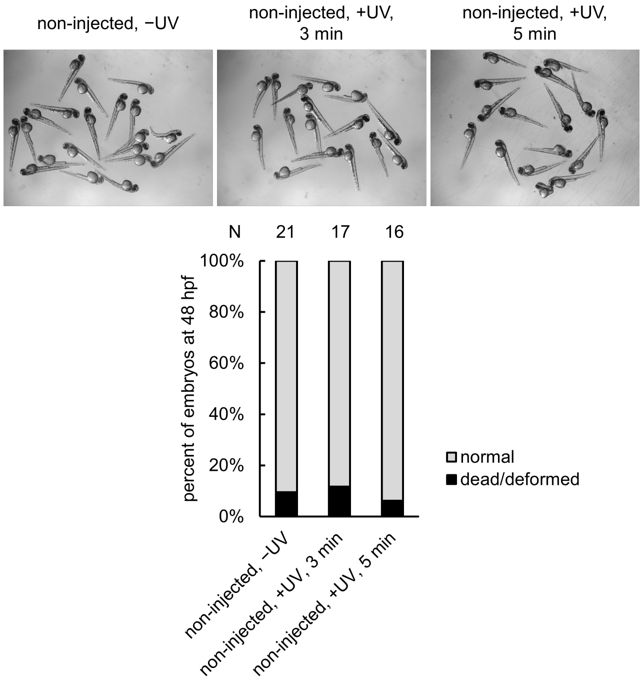

**Supporting Figure 12.** Non-injected embryos were irradiated with UV light as mentioned previously for 3 or 5 minutes at one hour post-fertilization. Embryos were imaged at 48 hpf and analyzed for toxicity. Micrographs of the fish are shown, with chart of those results below.

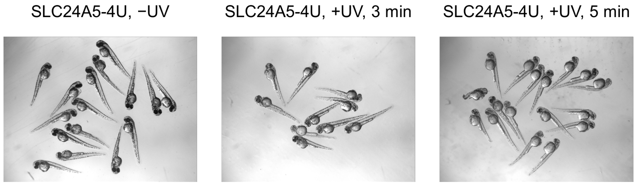

**Supporting Figure 13.** Micrographs of embryos injected with RNP targeting SLC24A5 and irradiated one hour after injection, then imaged at 48 hpf.
